# High-level carbapenem tolerance requires antibiotic-induced outer membrane modifications

**DOI:** 10.1101/2021.09.25.461800

**Authors:** Andrew N. Murtha, Misha I. Kazi, Richard D. Schargel, Trevor Cross, Conrad Fihn, Erin E. Carlson, Joseph M. Boll, Tobias Dörr

## Abstract

Antibiotic tolerance is an understudied potential contributor to antibiotic treatment failure and the emergence of multidrug-resistant bacteria. The molecular mechanisms governing tolerance remain poorly understood. A prominent type of β-lactam tolerance relies on the formation of cell wall-deficient spheroplasts, which maintain structural integrity via their outer membrane (OM), an asymmetric lipid bilayer consisting of phospholipids on the inner leaflet and a lipid-linked polysaccharide (lipopolysaccharide, LPS) enriched in the outer monolayer on the cell surface. How a membrane structure like LPS, with its reliance on mere electrostatic interactions to maintain stability, is capable of countering internal turgor pressure is unknown. Here, we have uncovered a novel role for the PhoPQ two-component system in tolerance to the β-lactam antibiotic meropenem in Enterobacterales. We found that PhoPQ is induced by meropenem treatment and promotes an increase in 4-amino-4-deoxy-L-aminoarabinose [L-Ara4N] modification of lipid A, the membrane anchor of LPS. L-Ara4N modifications enhance structural integrity, and consequently tolerance to meropenem, in several Enterobacterales species. Importantly, mutational inactivation of the negative PhoPQ regulatory element *mgrB* (commonly selected for during clinical therapy with the last-resort antibiotic colistin, an antimicrobial peptide [AMP]) results in dramatically enhanced tolerance, suggesting that AMPs can collaterally select for meropenem tolerance via stable overactivation of PhoPQ. Lastly, we identify histidine kinase inhibitors (including an FDA-approved drug) that inhibit PhoPQ-dependent LPS modifications and consequently potentiate meropenem to enhance killing of tolerant cells. In summary, our results suggest that PhoPQ-mediated LPS modifications play a significant role in stabilizing the OM, promoting survival when the primary integrity maintenance structure, the cell wall, is removed.

**Significance:** Treating an infection with an antibiotic often fails, resulting in a tremendous public health burden. One understudied likely reason for treatment failure is the development of “antibiotic tolerance”, the ability of bacteria to survive normally lethal exposure to an antibiotic. Here, we describe a molecular mechanism promoting tolerance. A bacterial stress sensor (PhoPQ) is activated in response to antibiotic (meropenem) treatment and consequently strengthens a bacterial protective “shell” to enhance survival. We also identify inhibitors of this mechanism, opening the door to developing compounds that help antibiotics work better against tolerant bacteria.

## Introduction

The rapid rise of antibiotic treatment failure threatens our ability to prevent and control bacterial infections. Antibiotic resistance, the continued proliferation of bacteria in the presence of the antibiotic, can often explain failure of clinical therapy. However, the response to an antibiotic is oftentimes more nuanced than a simple dichotomy of resistance vs. susceptibility. Bacteria can survive treatment in a non- or slowly-proliferating state, readily reverting to healthy growth after removal of the antibiotic (such as the end of a treatment course), and this is typically referred to as “antibiotic tolerance” (1–3). Importantly, tolerance to antibotics has been shown to enhance the evolution of outright resistance mechanisms (4–6), and can thus serve as both a direct and indirect contributor to treatment failure.

β-lactams are the most widely prescribed antibiotic class used to treat bacterial infections. The β-lactam ring inhibits the activity of the penicillin-binding proteins (PBPs) through covalent modification of a catalytic residue. PBPs are enzymes that synthesize the cell wall, an essential structure composed mainly of the polysaccharide peptidoglycan (PG). In many well-studied model organisms, PBP inhibition induces cell wall degradation and often subsequent lysis through the action of cell-wall degrading enzymes (collectively referred to as “autolysins”) in a poorly-understood manner (2). While lysis is the canonical response of model organisms like *Escherichia coli* K12, many susceptible clinical isolates of Gram-negative pathogens (including prominent Enterobacterales clinical isolates like *Klebsiella spp*. and *Enterobacter spp*.) exhibit a unique type of β-lactam tolerance. Like *E. coli*, these cells digest their PG upon exposure to β-lactams. However, instead of lysing, these pathogens survive antibiotic-induced cell wall degradation by forming viable, non-dividing, cell wall-deficient spheroplasts, which presumably rely on the outer membrane to counter their internal turgor (7–9). Interestingly, spheroplasts do not absolutely require osmotic stabilization and form in diverse types of growth media, including human serum (9). This cell wall-deficient phenotype is reminiscent of so-called L-forms (10–12), with the notable distinction that spheroplasts do not divide in the presence of the antibiotic.

Remarkably, spheroplasts formed in response to the carbapenem antibiotic meropenem readily resume growth and revert to wild-type rod shape when the β-lactam is removed from the growth medium (7, 9). Little is known about the molecular mechanisms that facilitate spheroplast formation and survival. In *Vibrio cholerae*, the two-component system (TCS) VxrAB is essential for spheroplast recovery by upregulating cell wall synthesis and downregulating iron uptake into the cells, mitigating toxic free iron levels induced by β-lactam treatment and allowing the cell to avoid damage by oxidative stress (13, 14). Many questions remain, however, as to how the cell envelope maintains its integrity without a cell wall, the essential structure canonically thought to protect the cell against immense turgor pressure.

In this study, we investigated genetic factors that contribute to the ability of *Enterobacter cloacae* to tolerate meropenem, which is used as a last-resort β-lactam to treat multidrug resistant bacterial infections (15–17). We first show that tolerance is dependent on outer membrane modifications (specifically 4-amino-4-deoxy-L-aminoarabinose [L-Ara4N]) induced by the PhoPQ TCS, an important cell envelope stress sensor that has previously been shown to respond to magnesium limitation, cationic antimicrobial peptide exposure, osmotic challenge, and pH changes (18). Both PhoPQ regulon transcription and resulting lipid A modifications are induced by meropenem treatment, suggesting a specific response to perturbations of PG synthesis in *E. cloacae*. These findings represent a novel mechanism of β-lactam tolerance in clinically relevant Enterobacterales, as well as an expanded role for the PhoPQ TCS.

## Results

### The PhoPQ TCS system regulates carbapenem tolerance

We previously showed that many Gram-negative pathogens are highly tolerant to meropenem. Upon treatment, tolerant cells do not appreciably lyse. Instead, they form viable, enlarged, non-replicating spheroplasts that are devoid of detectable cell wall material (9). Meropenem-induced spheroplast formation is quantifiable as an OD_600_ increase (**Fig. 1A**) and concomitant with only a moderate decrease in survival, as measured by colony-forming units (CFU) (**Fig. 1B**). In contrast, non-tolerant bacteria like many *E. coli* isolates rapidly lyse in the presence of meropenem, indicated by a decrease in both OD_600_ and survival (**Fig. 1AB**). Since spheroplast integrity is presumably maintained by the outer membrane, rather than the cell wall, we hypothesized that the strength of the outer membrane might correlate with tolerance. To test this, we repeated the killing experiments in the presence of the known outer membrane fortifying agents Mg^2+^ and Ca^2+^, which link adjacent lipopolysaccharide molecules by forming ionic bridges between phosphate groups on the lipid A domain (19, 20). Addition of either divalent cation (Mg^2+^ and Ca^2+^) supported spheroplast formation with a concomitant reduction in lysis during meropenem treatment in a concentration-dependent manner (**Fig. S1AB**). Furthermore, combinatorial addition of excess Ca^2+^ and Mg^2+^ completely prevented lysis (**Fig. S1A**). Thus, divalent cations prevent lysis during meropenem treatment, potentially through increasing the mechanical load-bearing capacity of the outer membrane through LPS crosslinking to protect the spheroplast structure.

**Figure 1:**
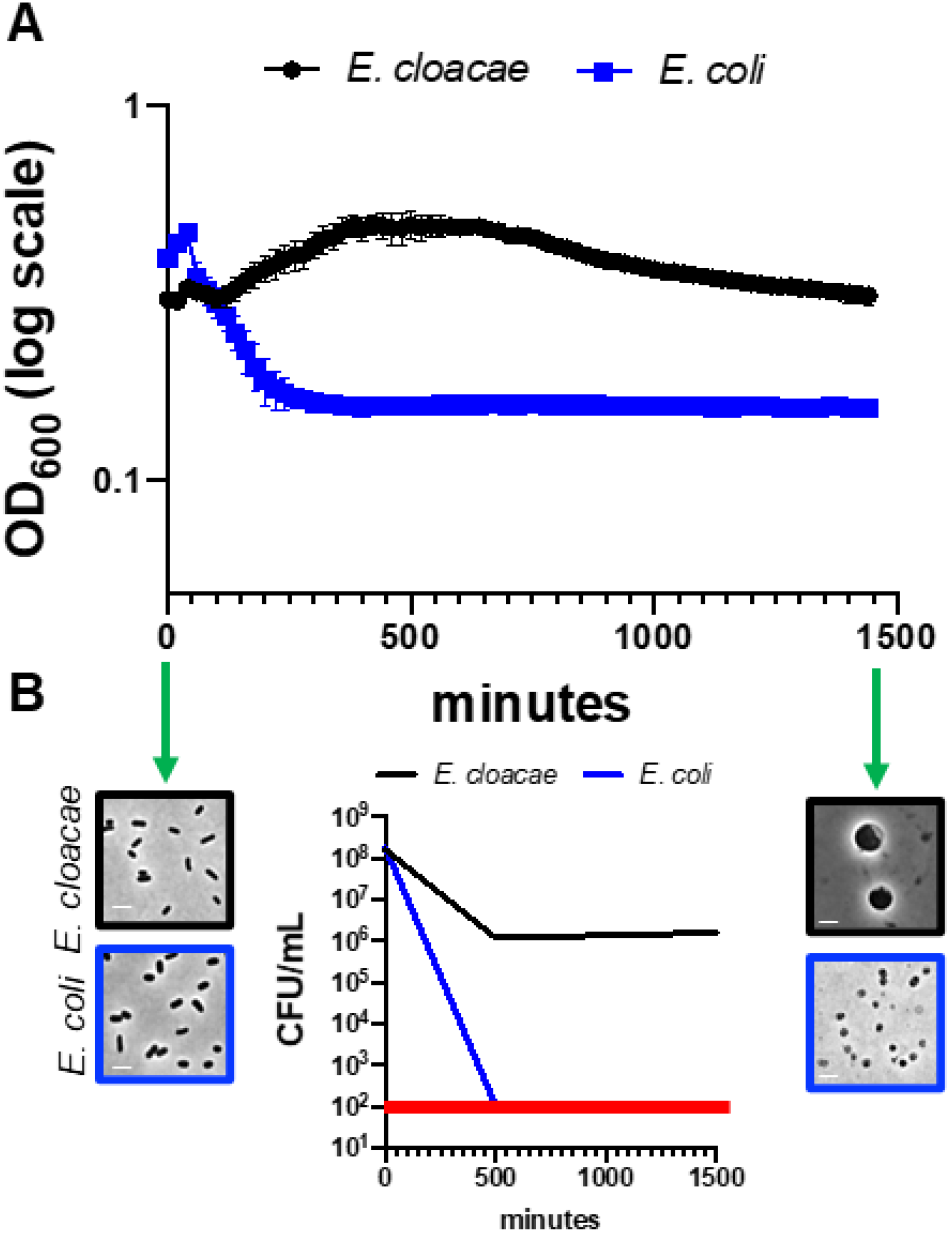
*Enterobacter cloacae* is highly meropenem tolerant. (A) Representative experiment demonstrating changes in OD_600_ measurements following meropenem treatment in *E. cloacae* relative to *E. coli*. Error bars represent the average of 3 technical replicates +/− standard deviation. (B) Survival was calculated as CFU/mL from the experiment depicted in (A). The red line denotes the limit of detection. Phase images from the same experiment show cells before and after meropenem exposure to illustrate spheroplast formation in *E. cloacae* and lysis in *E. coli* (only cell debris is visible in phase image). Scale bars, 2 μm.

When divalent cations are limiting, many Gram-negative pathogens induce outer membrane modifications (hyperacylation and/or increasing the positive charge of lipid A) to functionally substitute for divalent ionic bridges between lipopolysaccharide molecules (21). In *E. cloacae* (22) and other Enterobacterales (23), the PhoPQ TCS directly regulates expression of the *arn* operon, which synthesizes and transfers positively charged L-Ara4N moieties to lipid A, and *pagP*, which encodes an outer membrane acyltransferase (24). To test whether the PhoPQ TCS contributes to meropenem tolerance, we measured spheroplast formation in Δ*phoPQ*. Strikingly, OD_600_ declined sharply in Δ*phoPQ* relative to wild type, which could be fully complemented by ectopic expression of *phoPQ* (**Fig. 2A**). The decline in spheroplast formation strongly correlated with a robust 10-fold decrease in Δ*phoPQ* viability (CFU/mL) relative to wild type after 24 hours of treatment (**Fig. 2B**). Step-wise titration of Ca^2+^ and/or Mg^2+^ markedly enhanced Δ*phoPQ* tolerance (**Fig. S1CD**), suggesting decreased tolerance (spheroplast formation) in Δ*phoPQ* is due to its inability to crosslink adjacent LPS molecules in the outer membrane. To corroborate the involvement of PhoPQ, we sought to either enhance or reduce its activity and measured the effect of such perturbations on tolerance. PhoQ is antagonized by the small periplasmic MgrB protein; *mgrB* overexpression is thus expected to result in suppression of PhoPQ induction (25). Indeed, *mgrB* overexpression from a plasmid reduced spheroplast formation (proxied by OD_600_ measurements) (**Fig. 2C**) in wild type, closely resembling the Δ*phoPQ* phenotype (**Fig. 2A**). Conversely, exposing a strain deleted in *mgrB* to meropenem resulted in an increase in OD_600_ (**Fig. S2A**) that coincided with an approximately 1000-fold increased survival to meropenem relative to wild type (**Fig. 2B**). Modified lipid A structures in Δ*mgrB* were confirmed using matrix-assisted laser desorption ionization-time of flight mass spectrometry (MALDI-TOF MS) (**Fig. S3AB**). We also phenotypically validated our Δ*mgrB* mutant by assessing colistin MIC, which, as expected, was higher (≥128 μg/mL) than wild type (16 μg/mL). Collectively, our data suggest that the intensity of the PhoPQ response positively correlates with meropenem tolerance.

**Figure 2:**
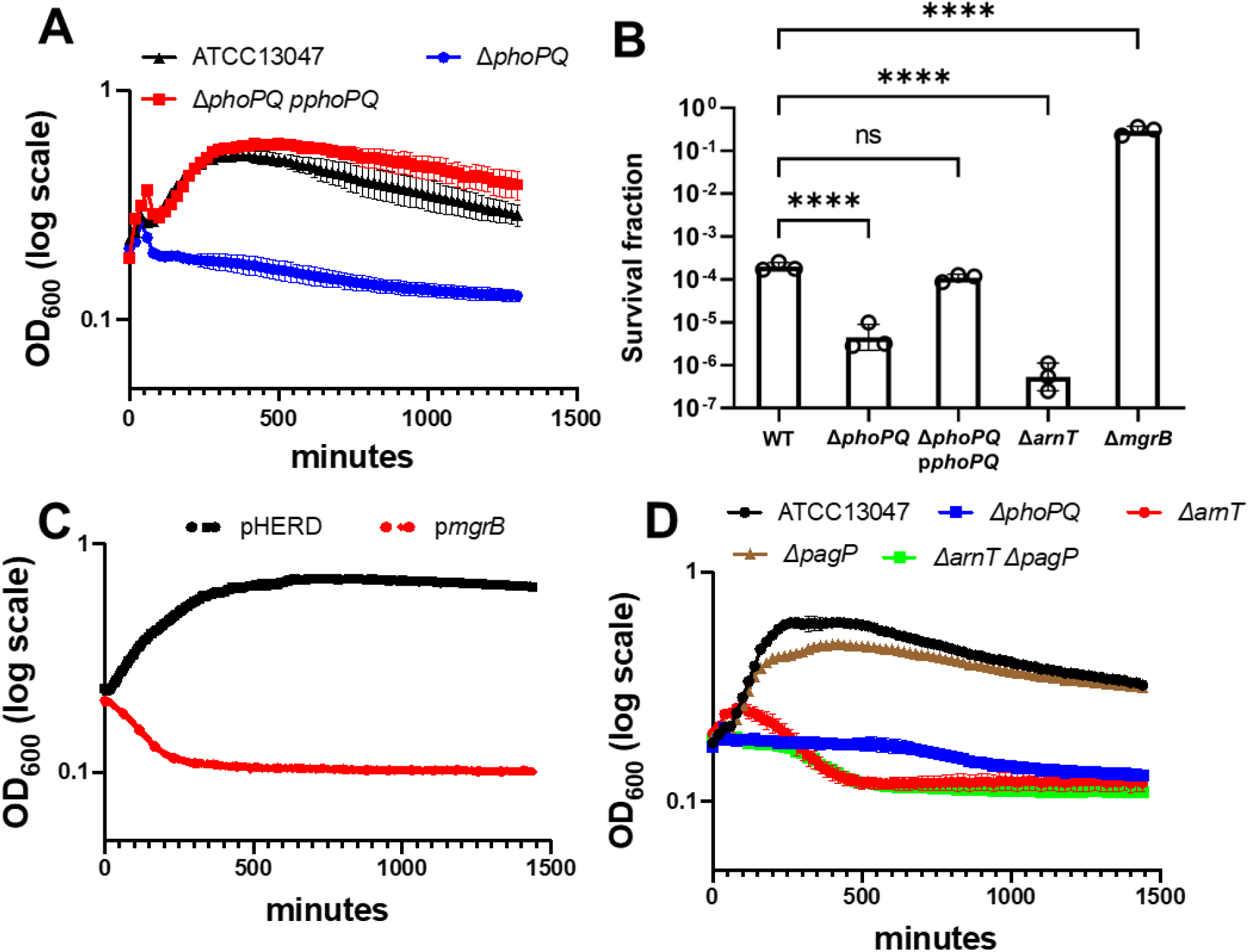
The PhoPQ system promotes meropenem tolerance in *E. cloacae*. (A) Spheroplast formation in response to meropenem treatment. Overnight cultures were diluted 10-fold into fresh LB medium containing 10 μg/mL meropenem and OD_600_ was measured. (B) Fraction of population surviving after 24 hours of meropenem exposure from experiments as described in (A). (C) MgrB overexpression reduces tolerance. Cells were treated as described in (A), but with the addition of 0.2 % arabinose (inducer). pHERD; empty vector. (D) The *arn* operon is required for meropenem tolerance. Experiments were conducted as described in (A). Data in each graph represent the average of 3 replicates +/− standard deviation; additional biological replicates are shown in Fig. S2B. Statistical significance for survival fraction determined by one-way ANOVA of log transformed data, followed by Tukey’s correction for multiple comparisons (ns, not significant; ****, *p* ≤ 0.0001).

To dissect individual contributions of PhoPQ-regulated genes to tolerance, we created mutants in the *pagP* and *arn* loci. While Δ*pagP* displayed spheroplast formation levels similar to wild type (**Fig. 2D**), Δ*arnT* exhibited a drop in OD_600_ reminiscent of Δ*phoPQ* (**Fig. 2D**) and a concomitant 100-fold decrease in CFU/mL relative to wild type (**Fig. 2B**). Notably, the survival defect was consistently more pronounced in Δ*arnT* vs. Δ*phoPQ* (**Fig. 2B**), suggesting either that residual L-Ara4N modification is retained in the absence of PhoPQ through basal expression of *arnT*, or that *phoPQ* induction in the absence of *arnT* is detrimental for an unknown reason. However, we observed notable variation in the extent of *arnT* tolerance measured by OD_600_; in some experiments, tolerance was reduced slightly compared to WT (**Fig. S2B**), while in others, the mutant approached Δ*phoPQ* levels (**Fig. 2D**). Thus, L-Ara4N modification appears to be the principal, but not the only, PhoPQ-regulated tolerance factor. Variability might reflect stochastic variation in another tolerance factor that at least partially mitigates the effects of *arnT* deletion.

### Meropenem induces *arn* transcription in a PhoPQ-dependent manner

Expression of the *arn* operon in *E. cloacae* is directly regulated by phosphorylated PhoP (22). Because we observed that PhoPQ was necessary for *arn-*mediated tolerance, we asked whether meropenem induced *arn* transcription in a PhoPQ-dependent manner. To test this, *E. cloacae* cells were exposed to meropenem for 30 minutes, after which *arnB*, *pagP*, and *phoP* transcript levels were quantified (**Fig. 3A**). Relative expression was calculated using 16s rRNA as an internal control. *phoP*, which is autoregulated (26), showed 2.5-fold higher expression in meropenem-treated cells relative to untreated. Additionally, *pagP* expression was 5-fold higher, and *arnB* expression was 6-fold higher after meropenem treatment. These results support a model where PhoPQ signaling, as well as transcription of its regulon, is induced in response to meropenem treatment.

**Figure 3:**
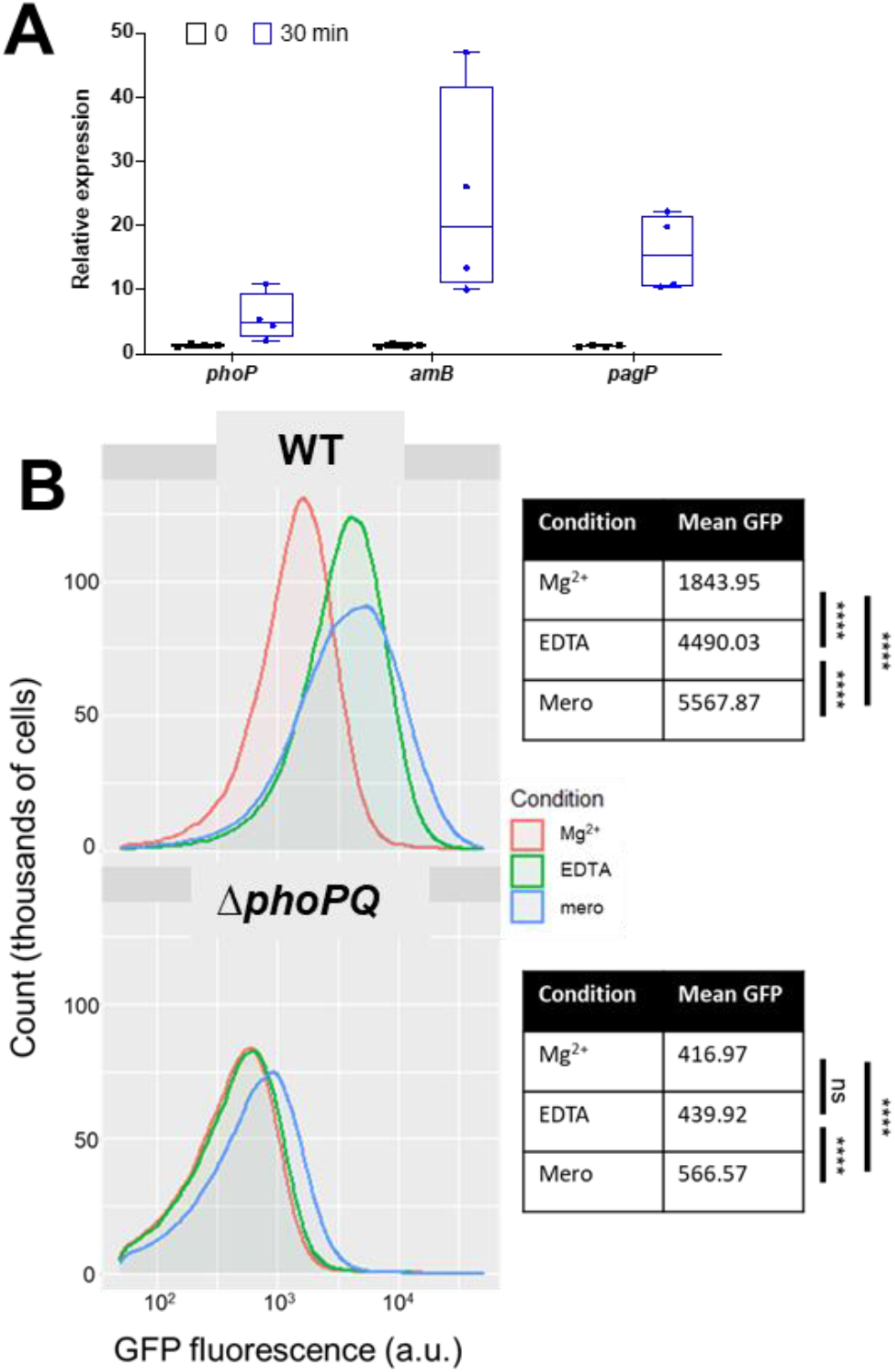
Expression of *arn*B in response to meropenem treatment is dependent on PhoPQ. (A) Relative expression reverse-transcription quantitative PCR (qRT-PCR) of PhoPQ regulon transcripts after meropenem exposure. Each experiment was independently replicated three times (individual data points of the three experiments are reported here). (B) Strains carrying transcriptional P_*arnB*_:msGFP fusions were exposed to indicated conditions and analyzed on a C6 Accuri flow cytometer. Statistical difference between populations was determined with a one-way ANOVA followed by Tukey’s correction for multiple comparisons (ns, not significant; ****, *p* ≤ 0.0001).

To corroborate the qRT PCR findings, we also constructed a fluorescent transcriptional reporter, fusing the *arnB* promoter (the first gene in the PhoPQ-regulated *arn* operon (22, 27)) with msfGFP, followed by fluorescence measurements upon exposure to meropenem (**Fig. 3B**). As a control, we first exposed cells to EDTA, which chelates divalent cations to destabilize the outer membrane and consequently activates PhoPQ (28). As expected, *ParnB*:msfGFP was induced by EDTA treatment in a *phoPQ*-dependent way (**Fig. 3B**). Interestingly, meropenem treatment also robustly activated the *ParnB*:msfGFP reporter, where a significant 3-fold fluorescence increase was measured, comparable to EDTA treatment. In contrast, meropenem only slightly (but reproducibly) increased *arnB* expression in the Δ*phoPQ* mutant under the same conditions. Thus, *arn* transcription is induced by meropenem in a primarily PhoPQ-dependent manner.

### Meropenem treatment promotes formation of L-Ara4N-modified lipid A species in a PhoPQ-dependent manner

The LPS lipid A domain is modified with L-Ara4N in a PhoPQ-dependent manner when Mg^2+^ is limiting (22), presumably to fortify the outer membrane during magnesium limitation stress. To determine if the *E. cloacae* lipid A structure is modified with L-Ara4N in response to meropenem treatment, we isolated lipid A from treated and untreated cultures. A distinct shift in lipid A structures was evident following 3 hours of meropenem treatment (**Fig. 4AB**), where increased ratios of L-Ara4N modified vs. unmodified forms were produced. Notably, the lipid A species that dominated before treatment (hexa-acylated, bis-phosphorylated, *m*/*z* = 1825.25) decreased in abundance in favor of Arn- and PagP-modified lipid A. Doubly Arn/PagP-modified lipid A (*m/z* = 2114.14) was also produced following treatment. Notably, while L-Ara4N modification of lipid A was PhoPQ-dependent, PagP-dependent lipid A acylation was not (see below for discussion). Quantitative thin-layer chromatography supported the MS results and revealed a 12.21 (+/− 1.13)-fold increase in single-modified L-Ara4-N lipid A in a PhoPQ-dependent manner (**Fig. 4C**).

**Figure 4:**
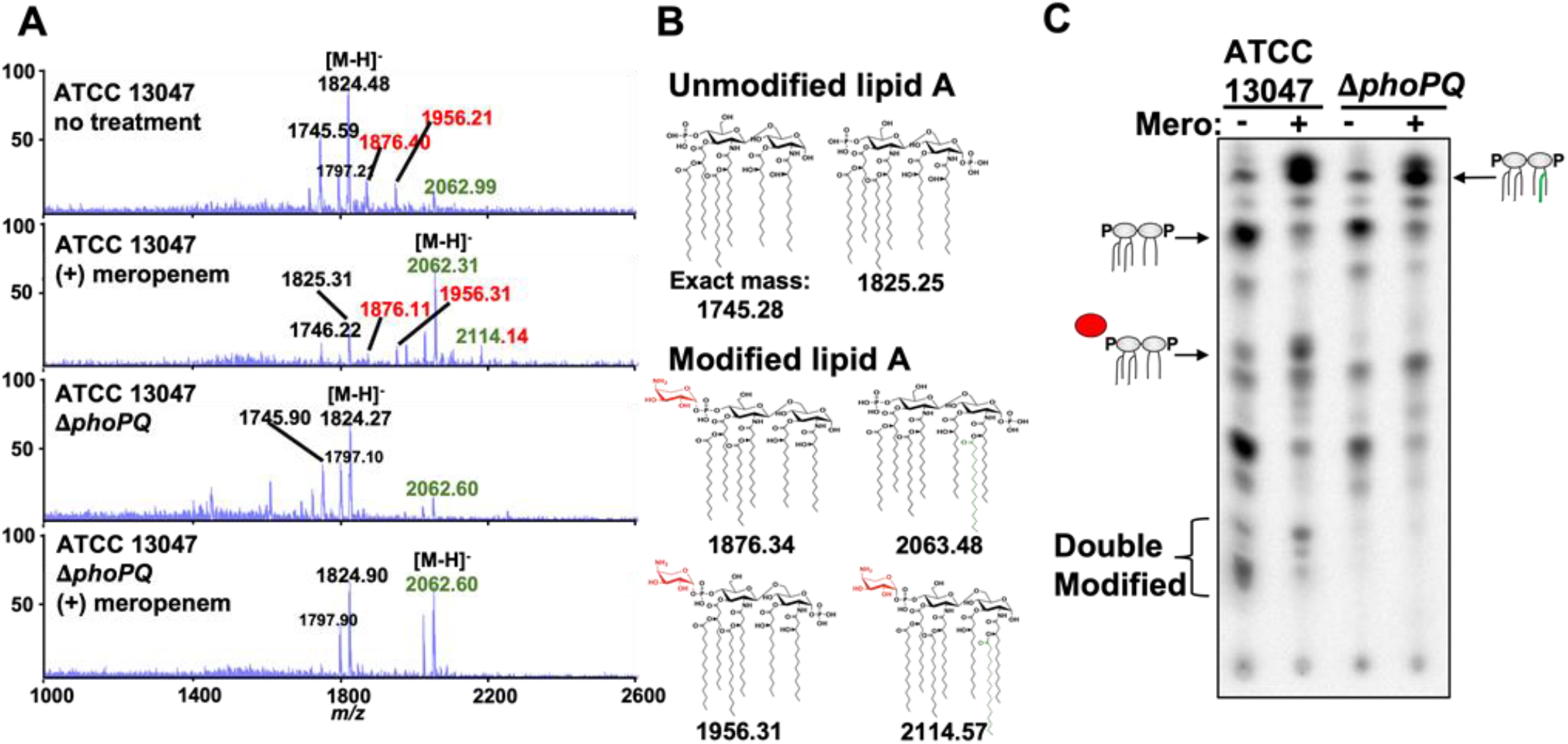
Analysis of *E. cloacae* lipid A after meropenem treatment. (A) MALDI-TOF MS analysis of lipid A extracted from wild-type or Δ*phoPQ E. cloacae* strains. L-Ara4N modifications are illustrated in red, while C_16:0_ additions are green. Numbered labels that are both red and green contain both modifications. Each experiment was independently replicated three times, and one representative data set was reported. (B) Predicted lipid A chemical structures in wild type and Δ*phoPQ E. cloacae*. (C) ^32^P-radiolabelled lipid A was extracted from treated or untreated wild-type or Δ*phoPQ E. cloacae*. Lipids were separated based on hydrophobicity using thin-layer chromatography. Red circle denotes L-Ara4N modification, while the green line indicates C_16:0_ addition.

*E. cloacae* encodes a principal putative PagP acyltransferase that we denoted as PagP1 (Ecl_03072) and PagP2 (Ecl_04468), a second apparent paralog. Lipid A extracted from Δ*pagP1* lacked the acyl chain induced upon meropenem treatment (**Fig. S4A**), suggesting that PagP1 removes palmitate (C_16:0_) from surface-exposed glycerophospholipids and transfers it to lipid A, as previously shown (29, 30), and demonstrating that PagP2 does not contribute to hyperacylation under these conditions. We also confirmed that meropenem-induced L-Ara4N modification was dependent on the *arn* operon (**Fig. S4B**). Furthermore, MS analysis of Δ*phoPQ* Δ*pagP1* (**Fig. S4C**) and Δ*arn* Δ*pagP1* (**Fig. S4D**) lipid A revealed that the mutants produced lipid A structures lacking all modifications following meropenem treatment, confirming that PagP1 and the *arn* operon products coordinate *E. cloacae* lipid A modifications in response to meropenem treatment. Interestingly, meropenem-induced hyperacylation was absent in Δ*mgrB* cells (**Fig. S3A**). We propose that meropenem-induced hyperacylation might actually be reflective of enhanced outer membrane glycerophospholipid bilayer accumulation in spheroplasts, a condition known for its likely ability to post-translationally activate PagP1 enzymatic activity (28, 31). This model is also consistent with our observation that while PagP1 is under genetic control of PhoPQ (22), meropenem-induced hyperacylation is independent of PhoPQ (**Fig. 4AC**). The absence of PagP1-dependent modification in Δ*mgrB* might indicate increased outer membrane strength (and concomitant reduction in glycerophospholipid in the outer leaflet of the outer membrane) in this background.

Based on the structural studies and our genetic evidence, we suggest PhoPQ-dependent tolerance is primarily mediated via L-Ara4N addition to lipid A. Presumably, L-Ara4N lipid A modification increases the structural integrity of the outer membrane through stabilization of lateral lipopolysaccharide interactions, which protects spheroplasts from internal turgor.

### Colistin exposure primes *E. cloacae* for meropenem tolerance

Cationic antimicrobial peptides (CAMPs) are known inducers of the PhoPQ TCS (32). Since our data above suggest that PhoPQ induction promotes tolerance, we hypothesized that pre-exposure to the CAMP colistin “primes” *E. cloacae* for tolerance to meropenem, either by inducing PhoPQ or by selecting for cells that have a higher baseline level of PhoPQ induction. To test this, we measured the extent to which *E. cloacae* was killed by meropenem with and without prior growth in colistin-containing media. Interestingly, after pre-exposure to colistin, the fraction of cells surviving meropenem treatment was approximately 10-fold greater (**Fig. 5**). This suggests that CAMPs have the potential to induce tolerance to other antibiotics, but that the temporal conditions of treatment may determine the extent to which this effect is significant.

**Figure 5:**
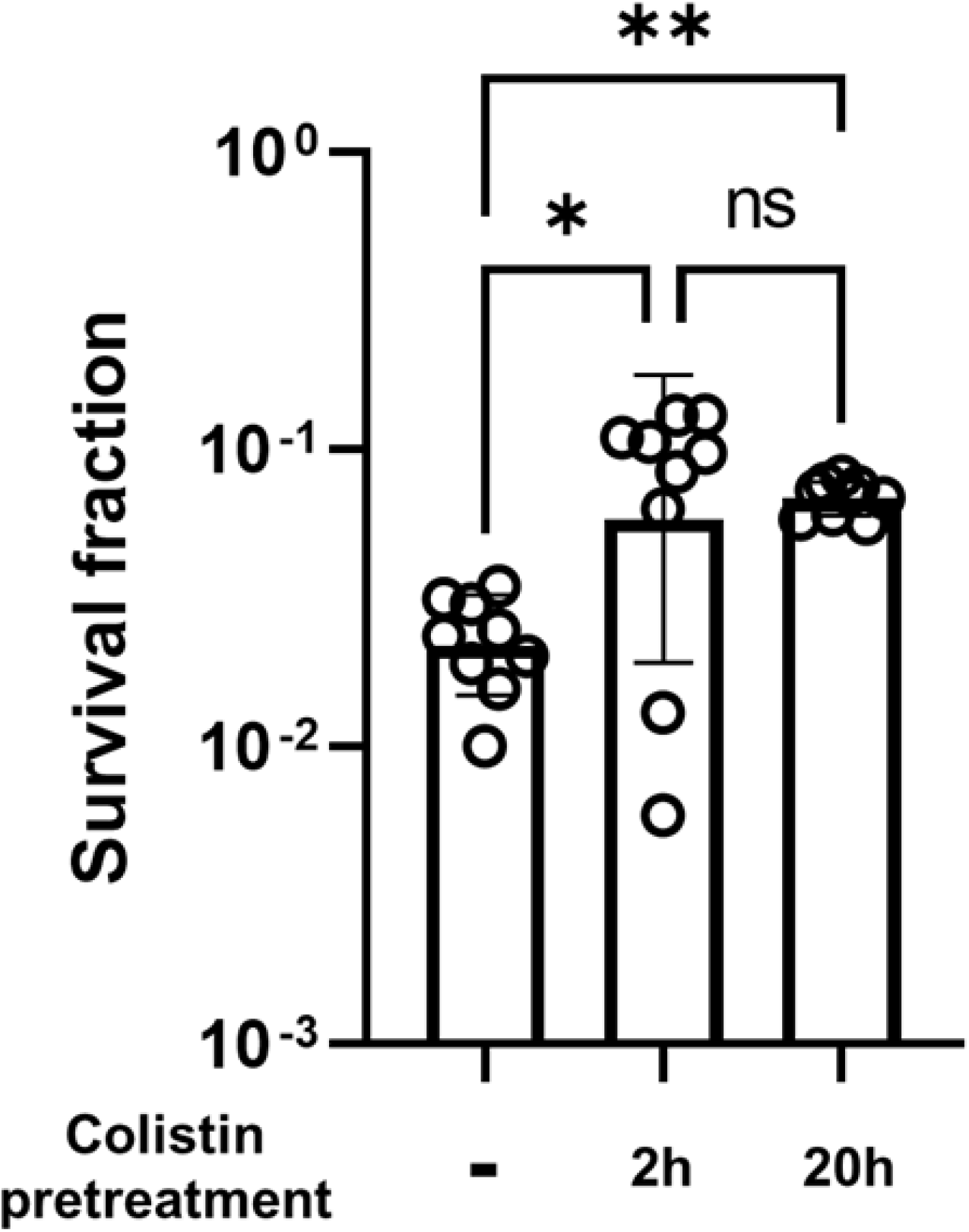
Colistin primes *E. cloacae* for meropenem tolerance. Overnight cultures were grown in LB +/− 20 μg/mL colistin, then diluted 10-fold into LB +/− 20 μg/mL colistin. Colistin pretreatment value represents total time exposed to colistin before exposure to 10 μg/mL meropenem. Cultures were then treated like other meropenem killing experiments (see methods). Survival fraction was calculated by dividing CFU/mL at 3h meropenem exposure by CFU/mL prior to meropenem exposure. Each bar represents the mean of 9 biological replicates, error bars represent standard deviation. Statistical significance determined by one-way ANOVA of log transformed data, followed by Tukey’s correction for multiple comparisons (ns, not significant; *, *p* ≤ 0.05; **, *p* ≤ 0.01).

### Outer membrane modifications are conserved β-lactam tolerance determinants in other Enterobacterales

We next sought to establish whether outer membrane modifications might promote tolerance in other Enterobacterales. We first turned to *Klebsiella pneumoniae* and used transposon insertion mutants in *phoP* and *phoQ* from the Manoil laboratory’s arrayed transposon mutant library (33) in killing experiments. After 6 hours of meropenem exposure, both the *phoQ::Tn* and *phoP::Tn* mutants exhibited 20- to 50-fold lower survival compared to the wild-type (**Fig. 6A**). We next used an *E. coli* K12 variant (WD101) engineered to constitutively upregulate the PmrAB two-component system (34) (which regulates L-Ara4N modification of lipid A in *E. coli*) to test the hypothesis that outer membrane modifications increase tolerance in *E. coli* (**Fig. 6B**). WD101 exhibited a dramatic, 10,000-fold increase in survival after 24 hours of meropenem exposure compared to the wild-type parental strain, further supporting a role for outer membrane modifications in meropenem tolerance.

**Figure 6:**
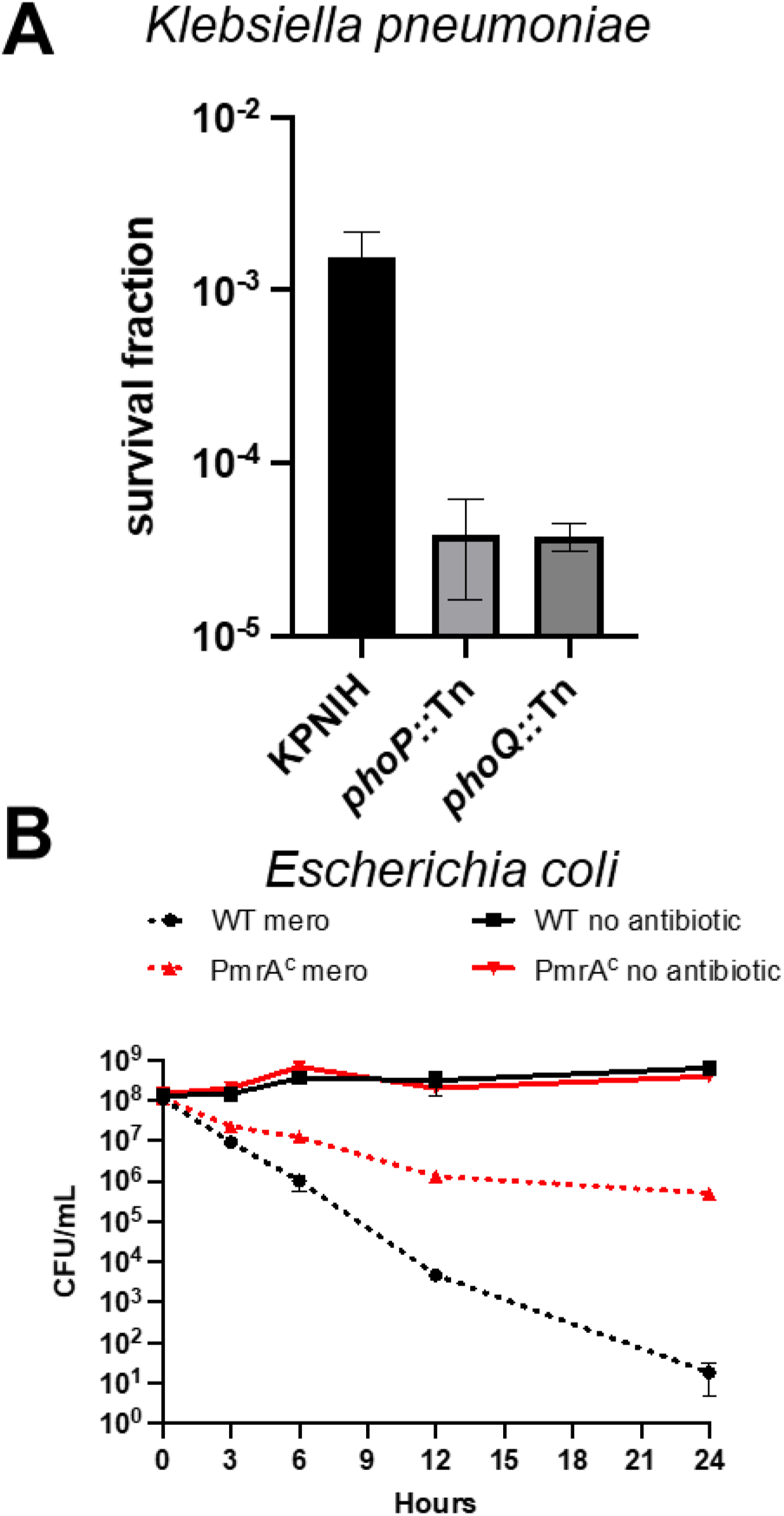
A conserved mechanism for meropenem tolerance in Enterobacterales. (A) *Klebsiella pneumoniae* KPNIH1 and its *phoP*::Tn and *phoQ::Tn* derivatives were diluted 10-fold into LB medium containing 10 μg/mL meropenem. Survival fraction is the CFU/mL after 6 hours of meropenem treatment divided by CFU/mL before treatment. Data represent averages of 6 biological replicates +/− standard error (B) *E. coli* strain W3110 (WT) or strain WD101, which has a constitutively active chromosomal copy of *pmrA* (*pmrA^C^*), were cultured in N-minimal medium and treated with or without meropenem. Cultures were incubated statically at 37°C. CFU were enumerated at 0, 3, 6, 12 and 24 hours. Error bars indicate standard deviation. Each experiment was independently replicated three times in triplicate, and one representative data set was reported.

### A small molecule inhibitor of PhoQ synergizes with meropenem and colistin in vitro

Tolerance is likely an under-appreciated contributor to antibiotic treatment failure. Antibiotic adjuvants that promote killing of tolerant cells thus have the potential to find a prominent place in our antibiotic armamentarium. Since histidine kinases like PhoQ are in principle targetable by small molecules, we tested whether his-kinase inhibitors synergized with meropenem. To this end, we turned to a previously developed suite of small molecules with potent histidine kinase inhibitory activity (35) and tested them in combination with meropenem. The anti-Amyotrophic Lateral Sclerosis (ALS) drug Riluzole, as well as its derivative Rilu-2, exhibited potent, concentration-dependent synergy in lysing tolerant *E. cloacae* cells *in vitro* (**Fig. 7A, Fig. S5A**). We also verified that Rilu-2 inhibited the formation of PhoPQ-dependent lipid A modifications using MALDI-TOF MS (**Fig. 7B**). Since the PhoPQ system is primarily recognized for its contribution to CAMP resistance in many Enterobacterales, we next tested the Rilu compounds’ ability to synergize with colistin. As expected, Rilu-2 and Riluzole indeed potentiated colistin-mediated killing (**Fig. S5B**), lending additional support to a PhoQ-inhibitory role of these compounds and also confirming previous results in *Salmonella* (36). Importantly, Riluzole is an FDA-approved treatment for ALS and could thus readily serve as an adjuvant against both meropenem-tolerant and colistin-resistant Enterobacterales.

**Figure 7:**
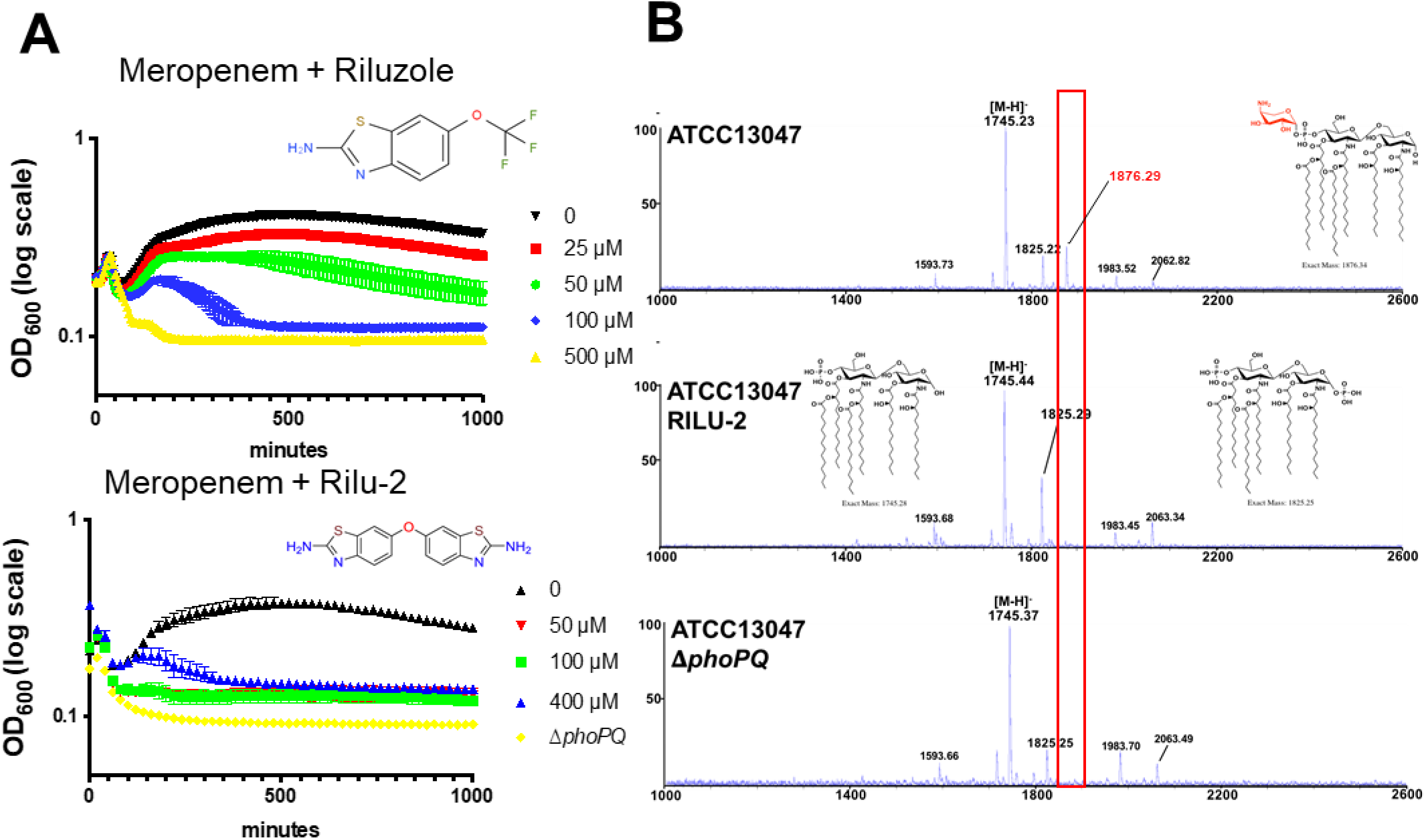
Rilu compounds synergize with meropenem to expedite *E. cloacae* killing. (A) Rilu compounds potentiate meropenem-induced lysis against *E. cloacae*. Overnight cultures were diluted 10-fold into fresh growth medium containing meropenem (10 μg/mL) and increasing concentrations of Riluzole or its derivative Rilu-2. Data represent the average of 3 technical replicates +/− standard deviation. (B) MALDI-MS analysis of lipid A isolated from untreated wild-type *E. cloacae*, cells treated with RILU-2 or Δ*phoPQ. m/z* corresponding with L-Ara4N modifications are illustrated in red. Relevant lipid A chemical structures are shown. Each experiment was independently replicated three times, and one representative data set is reported.

## Discussion

While much work has been done in Enterobacterales to elucidate mechanisms of antibiotic resistance and persistence, the genetic and molecular determinants of tolerance, and especially spheroplast formation, have remained poorly understood. In contrast to resistance (continued growth) and persistence (dormancy), carbapenem-tolerant populations are initially susceptible to treatment (i.e., lose their cell wall). This phenotype is reminiscent of so-called “L-forms” (10), with the notable difference that spheroplasts do not replicate in this state, while L-forms do. This is likely a consequence of L-forms being able to “escape” the outer membrane to then proliferate through stochastic membrane blebs (37, 38).

The remarkable ability of spheroplasts to survive without their PG layer lends support to the recent realization that the outer membrane has load-bearing capabilities (39) and prompted us to interrogate the molecular mechanism of outer membrane stabilization during antibiotic exposure. Our data suggest that L-Ara4N addition to lipid A is a key factor in spheroplast integrity. We propose that modifying lipid A molecules with a positive charge might increase outer membrane stability due to the enhancement of electrostatic interactions between adjacent LPS molecules on the surface-exposed face of the outer membrane. Of note, the L-Ara4N modification has been previously implicated in resistance to CAMPs, such as colistin (22). In this context, it is worrisome that colistin therapy can select for *mgrB* mutations (40–42), which we demonstrate here also confers high meropenem tolerance in addition to colistin resistance. Thus, treatment with antimicrobial peptide analogs (and, although we did not test this directly, potentially innate AMPs) can select for bacteria that are stably tolerant to subsequent therapy with a β-lactam. Our data also suggest that colistin (and, by extension, potentially AMPs of the innate immune system) can induce transient β-lactam tolerance, likely through induction of the PhoPQ system and perhaps other protective stress responses. However, the relationship between PhoPQ induction and tolerance is not absolute; a significant proportion of both colistin-pretreated and *mgrB-*deleted cells still lyse in the presence of meropenem. We speculate that β-lactam tolerance and colistin resistance, while relying on the same basic outer membrane modification, may each require a specific fraction of LPS molecules to be modified. Thus, within a sample, there may be a limited subset of cells exhibiting the correct amount of modification to enable both colistin resistance and meropenem tolerance (or even just optimal meropenem tolerance). This would explain why we observe only partial overlap between these two phenomena.

In summary, this work demonstrates a novel genetic determinant of carbapenem tolerance in clinically relevant Enterobacterales. Despite being a well-known regulator of polymyxin resistance, the PhoPQ two-component system was not previously known to respond or mediate tolerance to carbapenem treatment. As tolerance (and spheroplast formation in particular) is a possible culprit for antibiotic treatment failure (2, 3, 43), our results suggest a potential for combination therapies with histidine kinase inhibitors to increase the efficacy of carbapenems.

## Acknowledgements

Research on tolerance in the Dörr lab is supported by National Institutes of Health (NIH/NIAID) grant R01AI143704. Research in the Boll lab is supported by National Institutes of Health (NIH/NIGMS) grant GM143053. Histidine kinase inhibitor research in the Carlson lab supported by National Institutes of Health (NIH/NIGMS) grant GM134538-01A1 and the UMN Office of Academic Clinical Affairs.

## Materials and Methods

### Bacterial strains and growth

All strains/plasmids and primers used in this study are listed in **Table S1 and S2**. All strains were initially grown from freezer stocks on solid agar at 37°C. Isolated colonies were used to inoculate Luria-Bertani (LB), Brain heart infusion (BHI) or N-minimal medium at 37°C. Where required, kanamycin was used at 50 μg/mL, meropenem and colistin were used at 10 μg/mL, unless noted otherwise.

### Meropenem killing experiments

Unless noted otherwise, killing experiments were conducted in 100-well honeycomb plates in a Bioscreen C growth curve analyzer (Growth Curves USA, Piscataway NJ). Overnight cultures were diluted 10-fold into fresh LB medium containing meropenem (10 μg/mL, 300x MIC) and transferred to honeycomb plates (200 μL volume/culture). OD_600_ was measured by plate reader; at indicated timepoints, the experiment was paused and an aliquot was removed for CFU/mL determination or microscopy. Rilu compounds were dissolved in DMSO as 50 mM (Rilu-2) or 500 mM (Riluzole) stocks and added directly to the LB medium containing meropenem at the indicated concentrations.

### Colistin MIC experiment

Cultures were grown overnight at 37°C shaking, then diluted 1000-fold into fresh LB. Subcultures were grown for 1 hour at 37°C shaking before being diluted 1000-fold again into fresh LB to create a “seed culture”. 100 μL of seed culture was subsequently diluted 2-fold into a 96-well plate containing colistin concentrations ranging 0.25 – 128 μg/mL. Reported values are medians of 4 technical replicates.

### qRT-PCR

Relative-abundance quantitative PCR (qPCR) was performed as previously described (44, 45). In brief, the Sybr Fast One-Step qRT-PCR kit (Kapa Biosystems) was used with 16S rDNA as the internal reference. The PCR reaction was performed with Applied Systems RNA-Ct one-step system. Relative expression levels were calculated using the ΔΔCt method (46), with normalization of gene targets to16S rDNA signals.

### Flow cytometry GFP measurements

Cultures of strains harboring transcriptional *P_arnB_*:msfGFP fusions were grown overnight in LB supplemented with 10 mM MgSO_4_. Overnight cultures were then washed 2x in fresh LB before 10-fold dilution into fresh LB medium containing MgSO_4_ (10 mM),

Ethylenediaminetetraacetic acid (1 mM), or meropenem (10 μg/mL). Cultures were incubated statically for 3 hours at 37°C. Then, 500 μL of culture was harvested and run through a C6 Accuri flow cytometer (BD Biosciences) until 100,000 events (cells) had been analyzed. Mean green fluorescence as measured by the FL1-A channel was used as a readout for GFP.

### Mutant construction

*E. cloacae* subsp. *cloacae* 13047 mutant strains (*phoPQ*, *arn* and *arnT*) were constructed as previously described using recombineering with the plasmid pKOBEG (22, 47). Briefly, linear PCR products were amplified from pKD3 and transformed into *E. cloacae* ATCC 13047/pKOBEG strain by electroporation and plated on chloramphenicol selective media. Selected clones were transformed with pCP20 to cure the antibiotic resistance cassette. All mutants were verified by PCR.

*pagP1* was deleted using the Wanner method as described previously (22). Briefly, the chl resistance cassette was amplified from pKD3 using primers TDP1532/TDP1533, which contain 75 bp flanking homology overhangs. The resulting PCR product was electroporated into *E. cloacae* ATCC13047 expressing lambda red recombinase from pACBSR-hyg (48) (a hygromycin-resistant derivative of pKD46 (49)). Mutants were selected on chloramphenicol (100 μg/mL) and verified by PCR.

Other mutants were constructed using either lambda red recombinase (49) or the suicide vector pTox (50). The *mgrB* gene was deleted using the suicide plasmid pTox5 as described in (50). ~700 bp upstream and downstream flanking homology regions were amplified from ATCC13047 using primers TDP1767/68 and TDP1769/70, and cloned into pTox5 (digested with EcoRV) using isothermal assembly (51). Successful pTox5Δ*mgrB* were conjugated into ATCC13047 using the *E. coli* donor strain MFD lambda *pir*; successful recombinants were selected on plates containing 100 μg/mL chloramphenicol. Upon single colony purification, colonies were directly streaked out on an M9 minimal medium plate containing 0.2 % casamino acids and 1 % rhamnose, followed by incubation at 30 °C for 24 – 36 hours. Mutants were tested using primers TDP1771/72.

### Lipid A isolation and mass spectrometry

Isolation of lipid A for analysis was performed as previously described (52) with slight modifications. To analyze lipid A after meropenem treatment, overnight cultures grown in BHI broth were diluted 1:10 in pre-warmed media with or without meropenem statically for 3 h. To assess Rilu-dependent modification of lipid A, 12.5 mL of *E. cloacae* was grown to OD_600_ 1.0. Rilu-2 was used at a final concentration of 200 μM. Bacteria were harvested and lipid A extraction was carried out by mild-acid hydrolysis as previously described (53). For mass spectrometry (MS), data were collected on a MALDI-TOF (Axima Confidence, Shimadzu) mass spectrometer in the negative mode, as previously done (22).

For quantification of lipid A, cultures were grown with 2.5 μCi/mL of ^32^P orthophosphoric acid (^32^P) (Perkin Elmer) and lipid A was extracted. Thin layer chromotagraphy was done in a pyridine, chloroform, 88% formic acid, aqueous (50:50:16:5 v/v) tank for 3 hours. Plates were exposed to a phosphor screen, imaged, and densitometry was used to calculate the percentage of each lipid species. Reported densitometry was calculated using 2 replicates +/− standard deviation.

## Figure Legends

**Figure S1:**
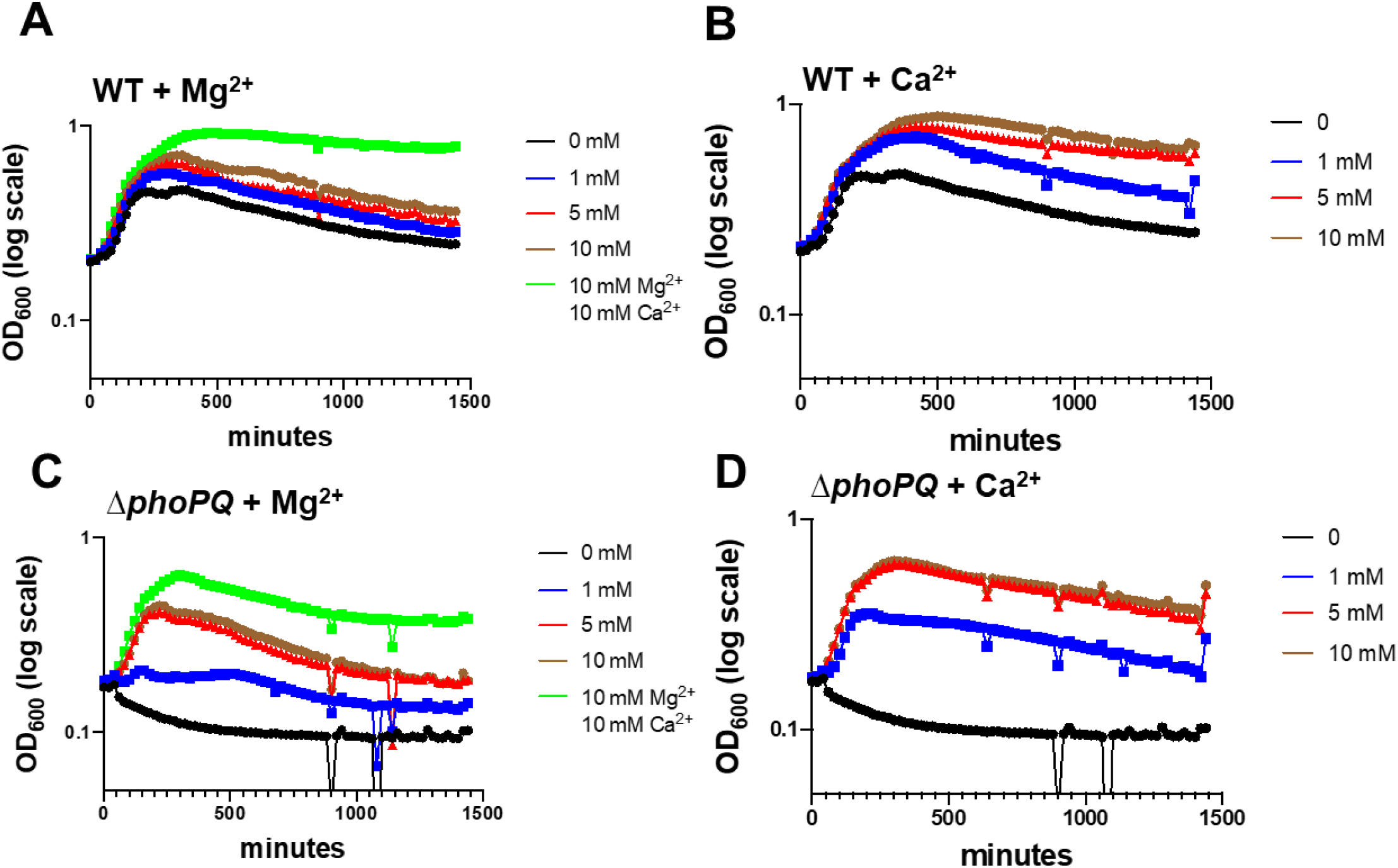
Addition of divalent cations prevents spheroplast lysis. Wild type (WT) (A-B) or its Δ*phoPQ* derivative (C-D) were treated as described in Fig. 1A with addition of the indicated concentrations of (A,C) MgSO_4_ (Mg^2+^) or (B,D) CaCl_2_ (Ca^2+^). Data represent the average of 3 replicates +/− standard deviation.

**Figure S2:**
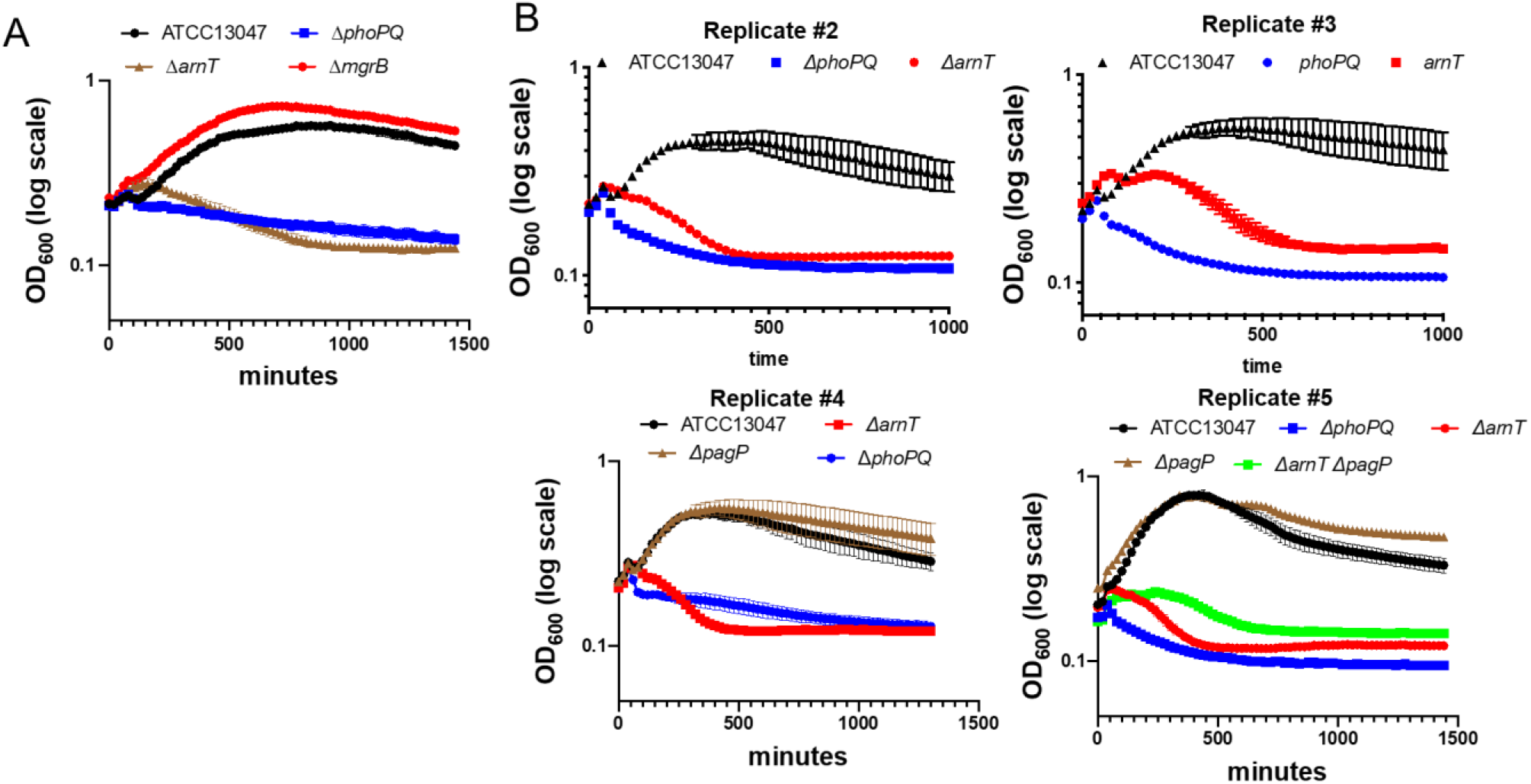
Independent biological replicates of experiments shown in Fig. 1. (A) An *mgrB* mutation promotes a moderate increase in mass increase during meropenem exposure. (B) Experiments were conducted as described in Fig. 1A legend; each graph represents experiments conducted on a different day. In addition, data in each graph represent the average of 3 biological replicates +/− standard deviation.

**Figure S3:**
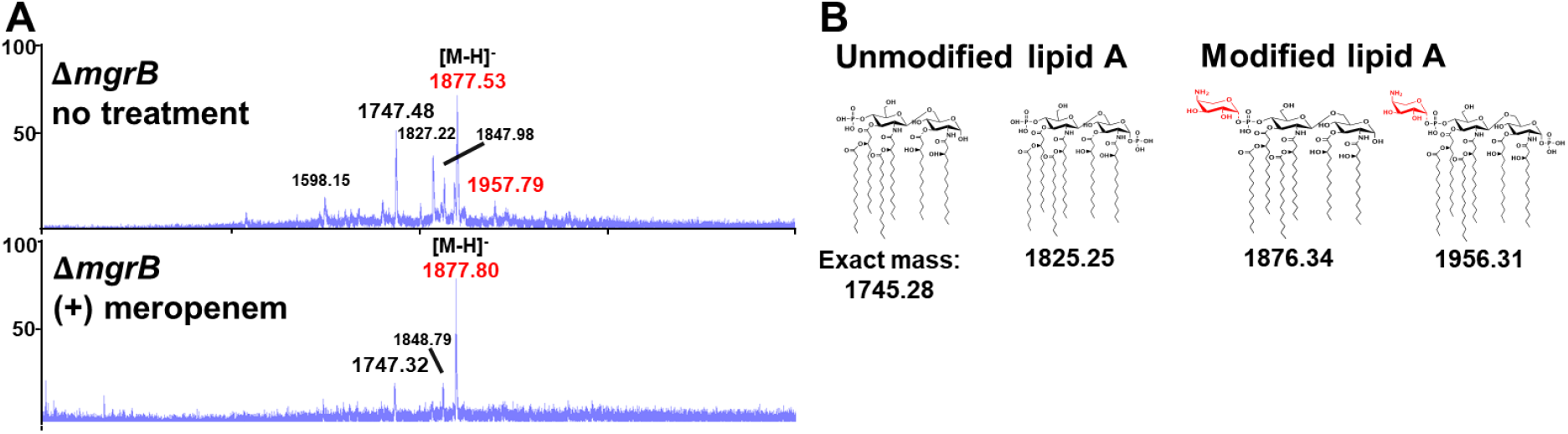
Analysis of lipid A from Δ*mgrB*. (A) MALDI-MS analysis of lipid A isolated from *E. cloacae* Δ*mgrB*. *m*/*z* corresponding with L-Ara4N modifications are illustrated in red. Each experiment was independently replicated three times, and one representative data set was reported. (B) Relevant lipid A chemical structures are shown.

**Figure S4:**
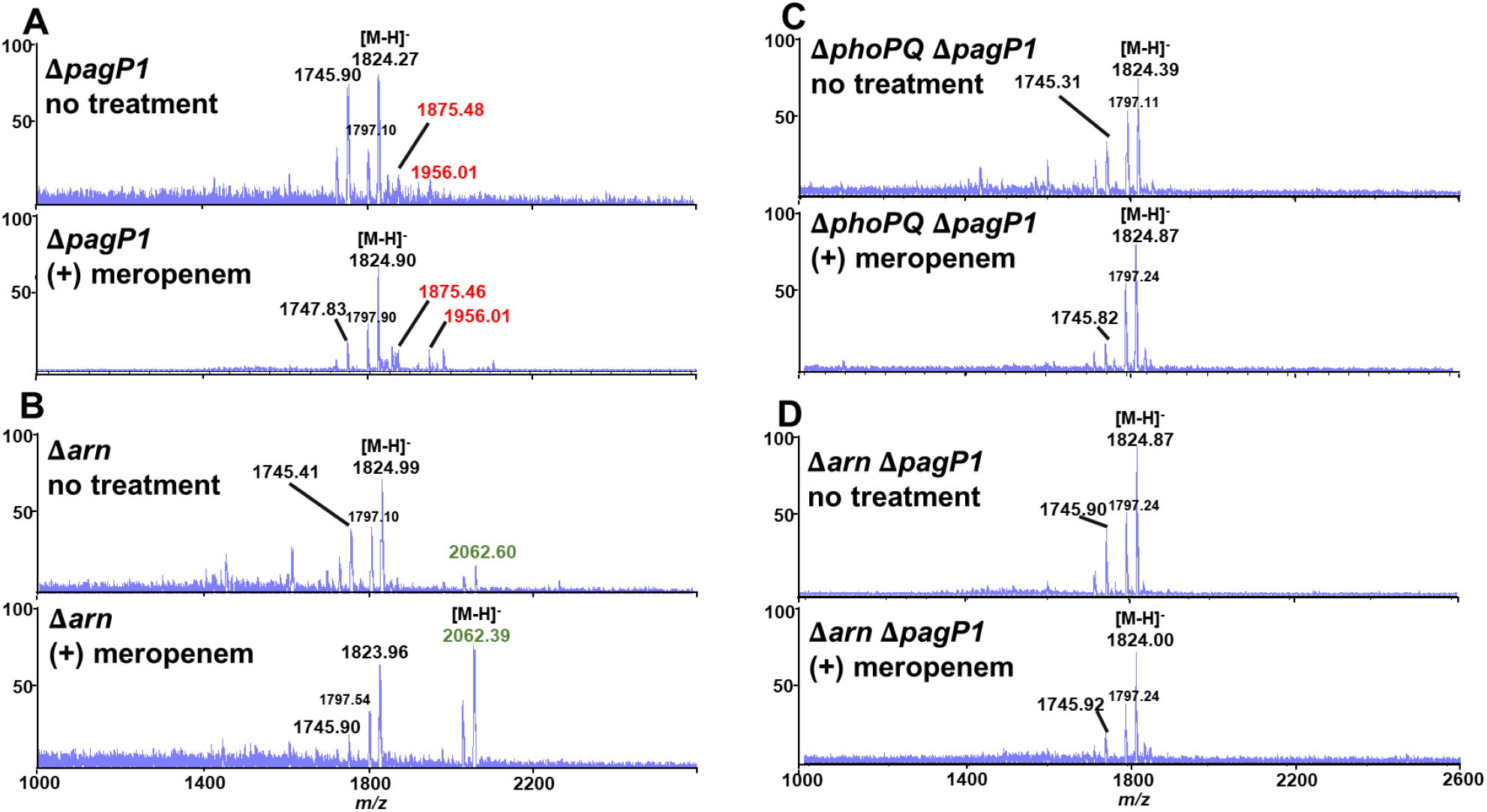
Analysis of lipid A from *E. cloacae* mutants. (A) MALDI-MS analysis of lipid A isolated from Δ*pagP1*, (B) Δ*arn*, (C) Δ*phoPQ* Δ*pagP1 and* (D) Δ*arn* Δ*pagP1*. *m*/*z* corresponding with L-Ara4N modifications are illustrated in red, while structures with altered acyl chain patterns are illustrated in green. Each experiment was independently replicated three times, and one representative data set was reported.

**Figure S5:**
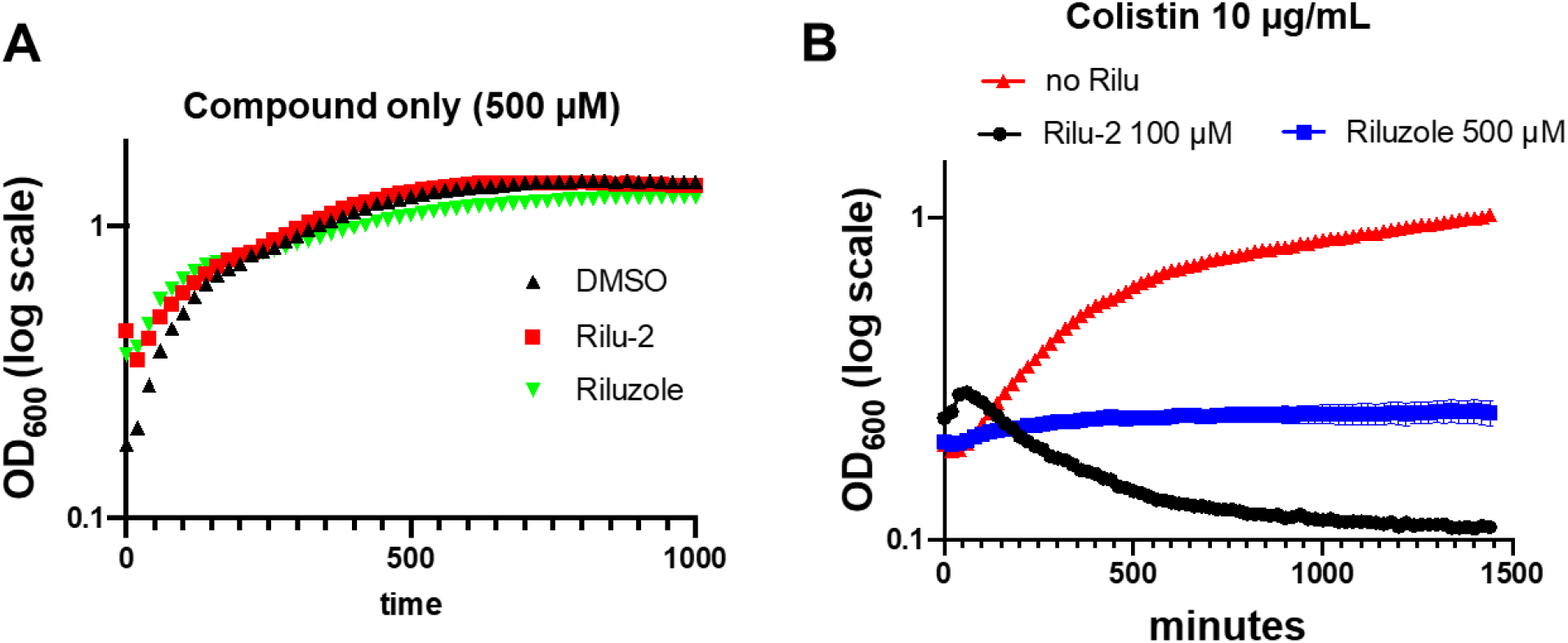
Rilu synergizes with colistin to enhance killing. (A) Rilu compounds do not cause lysis, but (B) potentiate colistin mediate killing. Experiments were conducted as described in Fig. 1A legend. Data represent the average of 3 replicates +/− standard deviation.

